# Reprogramming of host energy metabolism mediated by the TNF-iNOS-HIF-1α axis plays a key role in host resistance to *Plasmodium* infection

**DOI:** 10.1101/2024.03.26.586751

**Authors:** Kely C. Matteucci, Nathalia P. S. Leite, Patricia A. Assis, Isabella C. Hirako, Francielle Pioto, Ogooluwa Ojelabi, Juliana E. Toller-Kawahisa, Leonardo G. Vaz, Diego I. Costa, João S. Da Silva, José C. Alves-Filho, Ricardo T. Gazzinelli

**Author notes:** Contact info Gazzinelli, Ricardo T and Kely Catarine Matteucci.

## Abstract

TNF has a dual effect in *Plasmodium* infection, bolstering the host’s immune defense while also inducing sickness behavior. Here, we confirm that TNF signaling hampers physical activity, food intake, and energy expenditure while enhancing glucose uptake by the liver and spleen as well as controlling parasitemia in *P. chabaudi* (*Pc*)-infected mice. We also report that TNF is required for expression of inducible nitric oxide synthase (iNOS), stabilization of HIF-1α, expression of glucose transporter GLUT1 and enhanced glycolysis in monocytic cells from *Pc*-infected mice. Importantly, *Pc*-infected iNOS^-/-^, TNFR^ΔLyz2^ and *HIF-1α*^ΔLyz2^ mice show impaired release of TNF and glycolysis in monocytes, along with increased parasitemia and disease tolerance. Altogether, our results indicate that TNF-iNOS-HIF-1α-induced glycolysis in monocytes plays a critical role in host defense and sickness behavior in *Pc*-infected mice.

**Tease:** The role of host energy metabolism and glycolysis in monocytes as determinant of host resistance to *Plasmodium* infection and tolerance to disease.

## Introduction

Malaria is the most common mosquito-borne infectious disease caused by parasites from the *Plasmodium* genus (*1*). In 2021, approximately 247 million cases were reported, which resulted in nearly 619 000 deaths worldwide (*2*). The clinical symptoms include periodic fever episodes that occur synchronously with the rupture of infected red blood cells and the release of erythrocyte and parasite debris (*3*). Besides the characteristic development of fever, which is caused by systemic pro-inflammatory cytokine release (*3*), metabolic disorders like hypoglycemia and hyperlactatemia also occur in patients with malaria. High lactate plasma levels are related to the development of acidosis, which is an important prognostic factor in patients with severe malaria (*4–9*).

Successful immunity to *Plasmodium* infection requires the participation of monocytic cells, which have a central role in sensing and phagocytosing parasitized red blood cells (*10*). In experimental models, monocytes are the major cells responsible for secreting pro-inflammatory pyrogenic cytokines, such as IFN-γ, IL-1β and TNF, during acute malaria episodes (*11–13*). Among these pro-inflammatory cytokines, TNF plays an important role in the pathophysiology of malaria (*14*). In other contexts, TNF has been identified as a major inducer of glucose uptake and metabolism in host immune cells (*14–17*). Glucose is metabolized inside cells through glycolysis to generate energy and biosynthetic intermediates for cell growth, activation and proliferation (*18*, *19*). Moreover, the numerous intermediate metabolites that participate in the glycolysis-mediated conversion of glucose into pyruvate also play important roles in maintaining the activity and effector functions of innate immunity cells (*20*).

The hypoxia-inducible factor 1 alpha (HIF-1α) has emerged as one of the central regulators of inflammation and glucose metabolism in myeloid cells (*21*). Under normoxic conditions, HIF-1α is hydroxylated and degraded by the proteasome. Under hypoxic conditions, such as high levels of reactive nitrogen intermediates (RNI) and altered mitochondria function and release of reactive oxygen species (ROS), the activity of prolyl-hydroxylase domain (PHD) enzymes is inhibited, leading to stabilization of HIF-1α, and subsequent translocation to the nucleus (*22*, *23*). In addition, the expression of HIF-1α is upregulated in response to TNF. Consequently, HIF-1α activity in macrophages confers resistance to infection in different experimental models (*21*, *24–28*).

However, it is unclear how glucose metabolism, HIF-1α and their interplay with TNF affects innate immune cells and host resistance to *Plasmodium* parasites. Therefore, we investigated the involvement of TNF, RNI and HIF-1α in regulating glucose metabolism in monocytic cells and if this mechanism is relevant for host resistance and disease tolerance in experimental malaria. Our findings demonstrate that both TNF and inducible nitric oxide synthase (iNOS) promotes the expression and stabilization of HIF-1α enhance expression of glucose transporter GLUT1 and glycolysis in monocytic cells from *P. chabaudi*- (*Pc*-) infected mice. Altogether, our results reveal that this metabolic shift in monocytes has an important role in regulating the host energy metabolism, resistance to infection and disease outcome in an experimental malaria model.

## Results

### TNF signaling modulates signs of disease and energy expenditure in *P. chabaudi*-infected mice

TNF is involved in the development of protective immunity and clinical signs of malaria both in humans and experimental models (*29–31*). We first analyzed the time course of parasitemia in *Pc*-infected C57BL/6 mice and the blood glucose levels measured daily at the end of the dark cycle. Parasites were detected in the circulation, starting at 3 days post-infection (dpi), and the peak of parasitemia was observed at 8 dpi (Figure 1A). Blood glucose levels significantly decreased at 8 dpi, coinciding with the peak of parasitemia. At later time points, parasitemia decreased, and glycaemia returned to homeostatic levels (Figure 1B). As previously demonstrated (*32*), the circulating levels of TNF and expression of TNF mRNA in the liver peaked at 6 am (end of dark cycle) at 8 dpi (Figure 1C and 1D), and has been reported to peak between days 6 and 10 post-infection, with a consistent pattern observed on days 6 and 8 (*32*). We also observed that the percentage of iRBCs was higher in TNFR^-/-^ (TNFR p55p75^-/-^) compared to C57BL/6 mice (Figure 1E), as previously reported (*31*). In addition, while infected C57BL/6 mice displayed lower temperature in response to infection, the rectal temperature of infected and uninfected TNFR^-/-^ mice was similar (Figure 1F). Furthermore, the circulating levels of MCP1, TNF and IFN-γ were significantly lower in TNFR1^-/-^(TNFR p55^-/-^) compared to C57BL/6 mice (Figures 1G-I), whereas no difference was observed in IL-10 levels (Figure 1J). We next evaluated glucose homeostasis during *Pc* infection. Infected C57BL/6 mice exhibited increased glucose uptake, which was associated with the development of hypoglycemia at the peak of infection. In contrast, this metabolic alteration was not observed in TNFR^-/-^ mice, indicating that TNF signaling contributes to infection-induced changes in glucose metabolism and the subsequent development of hypoglycemia (Figure 1K). These results further indicate that TNF signaling promotes the systemic release of pro-inflammatory cytokine and plays a key role in controlling parasitemia, along with signs of disease in *Pc*-infected mice.

**Fig. 1.**
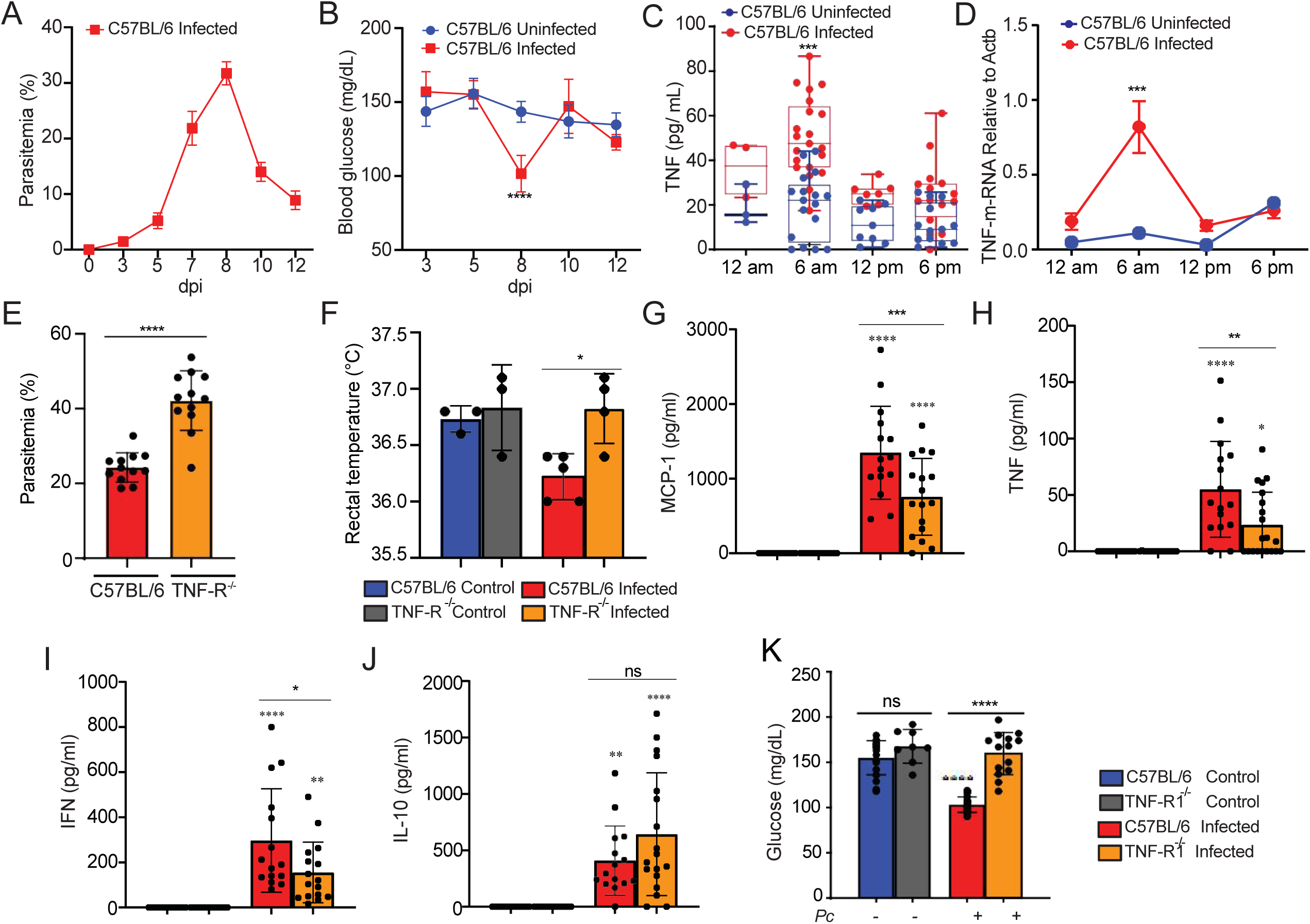
*P. chabaudi*-infected mice display enhanced TNF production associated with increased disease parameters. C57BL/6 mice were infected or not i.p. with *P. chabaudi* (10^5^ iRBCs). **(A)** Parasitemia was determined at 0, 3, 5, 7, 8, 10 and 12 days-post infection (dpi). **(B)** Blood glucose levels were measured with a glucometer at 0, 3, 5, 7, 8, 10 and 12 dpi. Means of blood glucose levels were compared between infected (red) and uninfected (blue) mice at the same time points. **(C)** Blood TNF Levels were measured by ELISA at 8 dpi. Means of blood TNF levels were compared between infected (red) and uninfected (blue) mice at the same time points. **(D)** TNF m-RNA from mice liver was determined at 8 days post-infection (dpi). **(E)** Parasitemia was determined at 8 days-post-infection (dpi) in C57BL/6 and TNFR deficient mice. **(F)** Rectal temperature was determined at 8 dpi in C57BL/6 and TNFR deficient mice. **(G-J)** MCP-1, TNF, IFN and IL-10 was measured by CBA Kit at 8 dpi. **(K)** Blood glucose levels were measured with a glucometer at 8dpi. A, B and F: Graphs show mean ± SEM of 1 representative experiment of at least 3 independent ones performed with 4-5 mice per group; Statistical analysis: One-way ANOVA with Tukey post-hoc test. C and D: Graph shows mean± SEM of combined data from 3 independent experiments with 4 - 6 mice per group; Statistical analysis: One-way ANOVA with Bonferroni post-hoc test. E, G, H, I, J, K: Graphs show mean ± SEM of combined data from 3 independent experiments with 4-6 mice per group; Statistical analysis: One-way ANOVA with Tukey post-hoc test. Asterisks above the bars indicate comparisons between mice of the same strain before infection and at 8 dpi. Asterisks connected by horizontal lines indicate comparisons between infected groups. *P≤0.05, **P≤0.01, ***P≤0.001, **** P≤0.0001, ns = non-significant.

To further address the role of TNF signaling in promoting disease signs in *Pc*-infected mice, we evaluated alteration on host energy metabolism, as indicated in physical activity, food intake, overall energy expenditure and respiratory exchange rate in naïve or *Pc*-infected C57BL/6 and TNFR^-/-^ mice during 24h, at the 8 dpi. We observed that in naïve animals, all these parameters were similar in TNFR^-/-^ and C57BL/6 mice (Figures 2A-D, top panels and Figures 2E-H). In contrast, all the evaluated parameters were decreased in infected C57BL/6 mice compared to their naïve counterparts during the light and dark cycles. When we analyzed only infected mice, the alterations in all parameters were milder in TNFR^-/-^compared to C57BL/6 mice (Figures 2A-D bottom panels and 2E-H).

**Fig. 2.**
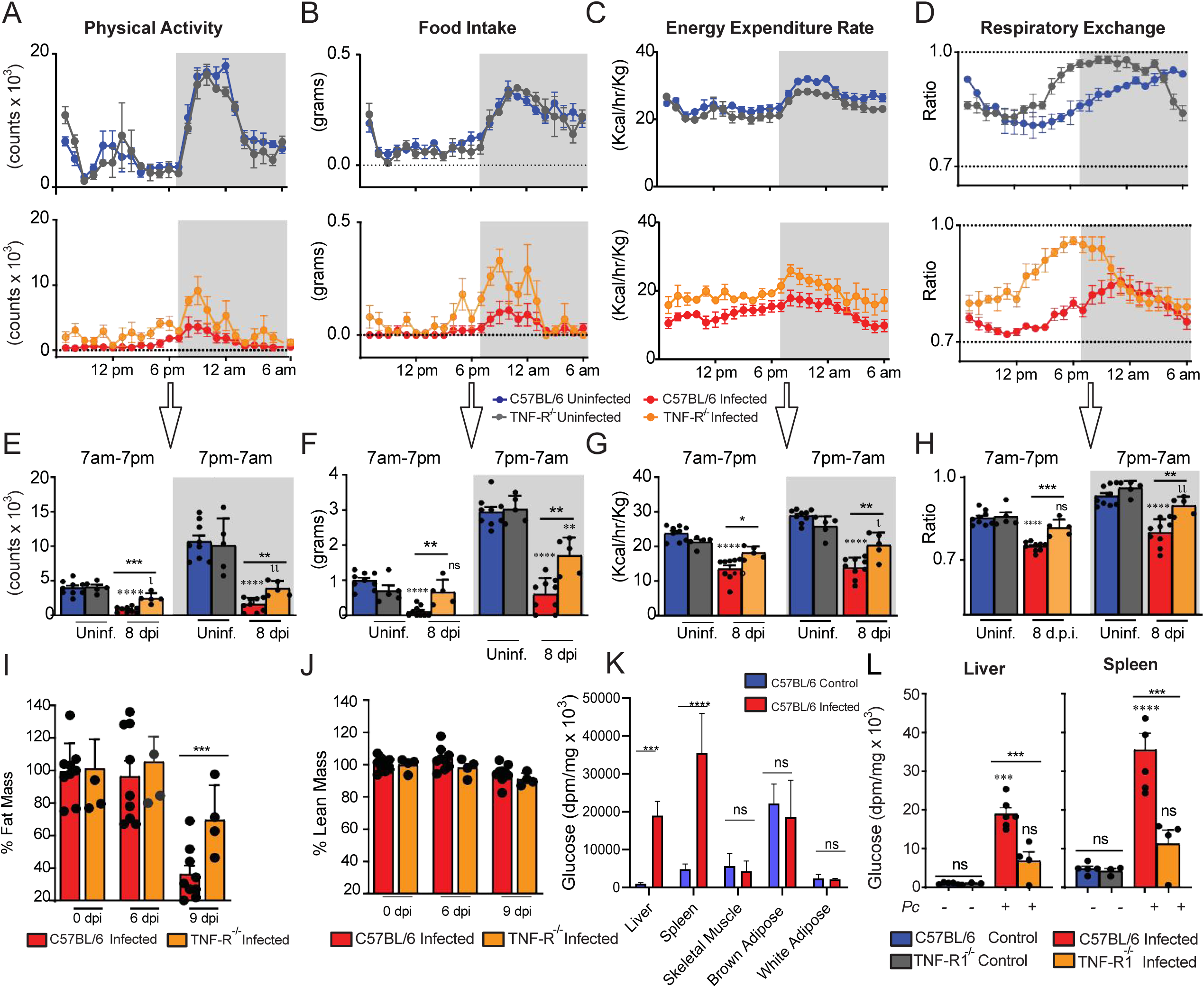
The detrimental effects of *P. chabaudi* infection on physical activity, food intake and energy metabolism are partially dependent on TNF receptor signaling. Mouse energy balance from C57BL/6 and TNFR^-/-^ was assessed using metabolic cages (TSE 959 Systems, Chesterfield, MO) at 3 days prior to infection (uninfected) (C57BL/6: blue, TNFR-/-: gray) and at 8 days postinfection (dpi) with P. chabaudi (10^5^ infected RBCs) (C57BL/6: red, TNFR^-/-^: orange). Scores for **(A and E)** physical activity, **(B and F)** food intake, **(C and G)** energy expenditure, **(D and H)** respiratory exchange, and **(I and J)** percentual of fat mass and lean mass are depicted. Hourly measurements of such parameters are shown in top row panels for mice 3 days prior to infection (uninfected) and in middle row panels for infected mice. The average values for light and dark cycles in all groups are depicted in bottom row panels. **(K and L)** Glucose uptake was measured 2-[14C] deoxyglucose. Experiments were performed at the National Mouse Metabolic Phenotyping Center (MMPC) at UMASS Medical School. Basal glucose uptake in individual organs of infected (red) and uninfected (blue) mice was measured using an intravenous injection of 2-[14C] deoxyglucose. After 1h, mice were anesthetized, and tissue samples were taken for organ-specific levels of 2-[14C] deoxyglucose-6-phosphate. A - L: Graph shows mean ± SEM of combined data from 2 independent experiments; Statistical analysis: One-way ANOVA with Tukey post-hoc test. Asterisks above the bars indicate comparisons between mice of the same strain before infection and at 8 dpi. Asterisks connected by horizontal lines indicate comparisons between infected groups. *P≤0.05, **P≤0.01, ***P≤0.001, **** P≤0.0001, ns = non-significant.

We then asked whether *Pc* infection might also affect body fat mass and tissue glucose uptake. We observed a major depletion of fat mass in infected C57BL/6 mice at 9 dpi, which was less pronounced in TNFR^-/-^ animals (Figure 2I), while the percentage of lean mass was not altered in either wild type or TNFR^-/-^mice (Figure 2J). We also found that *Pc* infection resulted in a marked increase in glucose uptake in the liver and spleen of C57BL/6 mice, unlike skeletal muscle or white and brown adipose tissue (Figure 2K). The increase in these organs was not seen in TNFR^-/-^ animals (Figure 2L). Altogether, these results demonstrate the role of TNF in regulating glucose and energy metabolism in *Pc*-infected mice and, consequently, with important implications in the pathophysiology of experimental malaria.

### Hepatic non-parenchymal cells display enhanced GLUT1 receptor expression and shift to glycolytic metabolism following *P. chabaudi* infection

The liver has a key role in regulating host glucose metabolism (*33*). Therefore, we asked whether *Pc* infection alters the expression of genes related to carbohydrate metabolism in the liver. There was an overall enhancement in the expression of genes related to glycolysis and a reduction in the expression of genes associated with the Tricarboxylic (TCA) cycle and gluconeogenesis (Figure 3A). Enhanced glycolysis relies on the heightened glucose uptake by cells, mediated by different glucose carriers (*34*, *35*). However, the expression of GLUT2 (*Slc2a2),* GLUT5 *(Slc2a5)* and GLUT9 *(Slc2a9)* was decreased, whereas of GLUT1 (*Slc2a1),* GLUT3 (*Slc2a3)* and GLUT6 *(Slc2a6)* expression was increased in the liver of C57BL/6 infected, as compared to naïve mice (Figure 3A). Among the transporters whose expression was increased, GLUT3 is expressed mainly in neurons (*36*), while *Slc2a6* GLUT6 has been shown to be located in lysosomes and does not mediate glucose uptake (*37*). GLUT1, on the other hand, is a ubiquitously expressed and highly effective transporter of glucose. Also, GLUT1 is the most well-characterized surface glucose transporter in immune cells (*18*, *38*, *39*). Therefore, we next quantified GLUT1 protein, and, as expected, an enhancement in GLUT1 levels was found in the liver of *Pc*-infected C57BL/6 mice, as compared to the uninfected controls (Figure 3B).

**Fig. 3.**
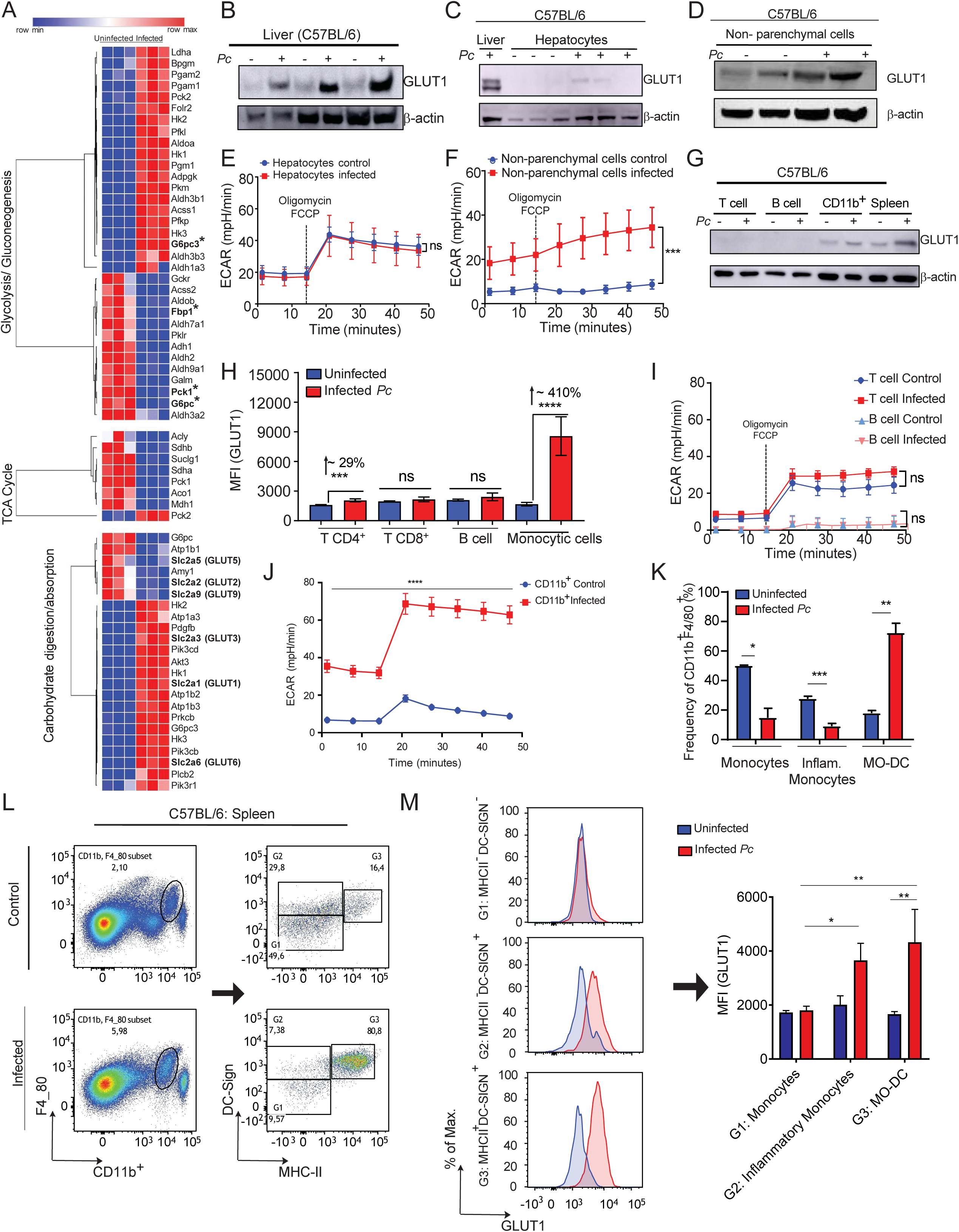
*P. chabaudi* infection stimulates Increased GLUT1 expression in hepatic non-parenchymal and splenic CD11b+ cells associated with an altered host metabolic profile. C57BL/6 mice were infected or not i.p. with P. chabaudi (10^5^ iRBCs). **(A)** RNA-seq was performed in livers from infected and uninfected mice. **(B)** GLUT1 expression in livers of infected (8 dpi) or uninfected mice was evaluated on by western blot. **(C and D)** GLUT1 expression in **(C)** hepatocytes or **(D)** hepatic non-parenchymal cells isolated from livers from C57BL/6 infected (8dpi) or uninfected mice, quantified by western blot (expression of β-actin was used as control). **(E)** Hepatocytes or **(F)** Hepatic non-parenchymal cells were purified from livers harvested from infected (red - 8 dpi) and uninfected (blue) mice and cultured ex vivo for quantification of extracellular acidification rate (ECAR) by Seahorse XFe96. **(G, I and J)** T cells, B cells and myeloid cells (CD11b+) were purified from splenocytes of infected and uninfected mice using magnetic beads and **(G)** GLUT1 expression was evaluated by western blot. **(I)** T and B cells or **(J)** CD11b+ purified from spleens were cultured *ex vivo* for quantification ECAR. **(H, K, L and M)** graphs showing GLUT1 expression evaluated by flow cytometry in splenic cells from infected (red – 8 dpi) and uninfected (blue) mice. A. Heatmap represents 3 biological replicates performed in RNA-seq. B, C, D and G: Western blot images representative of at least 3 independent experiments. E, F, I and J: Graphs show mean ± SEM of 1 representative experiment of at least 3 independent ones performed with 4-5 mice per group; Statistical analysis: Two-way ANOVA followed by Sidak’s post-hoc test employing mixed effect analysis provided by Agilent Seahorse XFe96 analyzer software. H, K L and M: Representative dot plots and Graphs show mean ± SEM of 1 representative experiment of at least 3 independent ones performed with 4-5 mice per group; Statistical analysis: Student’s t test. Asterisks above the bars indicate comparisons between mice of the same strain before infection and at 8 dpi. Asterisks connected by horizontal lines indicate comparisons between infected groups. *P≤0.05, **P≤0.01, ***P≤0.001, **** P≤0.0001, ns = non-significant.

Hepatocytes have an important role in glucose uptake from the circulation, and they do this majorly through GLUT2 (*38*), whose mRNA expression was downregulated (Figure 3A). Since GLUT1 expression can also be induced in hepatocytes (*18*), we evaluated if GLUT1 expression was enhanced in hepatocytes and non-parenchymal cells from the liver of infected mice. We found that at 8 dpi, GLUT1 expression was not altered in hepatocytes (Figure 3C), but increased in non-parenchymal cells (Figure 3D). We have also assessed the metabolic activity of these same hepatic cell populations by quantifying their extracellular acidification rate (ECAR) and oxygen consumption rate (OCR), which are indicatives of how much glucose is being metabolized through glycolysis and the TCA cycle, respectively. Consistent with GLUT1 expression, we observed no change in the metabolic profile of hepatocytes (Figure 3E) but an increase in ECAR of non-parenchymal cells (Figure 3F) from the liver of *Pc*-infected mice.

### CD11b^+^ cells from the spleen shift towards glycolytic metabolism in response to *P. chabaudi* infection

The spleen is the main tissue involved in immunity to *Plasmodium* infection, where the response of immune cells to infection results in remarkable splenomegaly (*40*). Moreover, the magnitude of glucose uptake in spleens was higher than in the livers of infected mice (Figure 2K). Hence, we next asked whether the increased tissue glucose uptake in experimental malaria was associated with the enhancement of GLUT1 expression and glycolysis in different subsets of immune cells. The levels of GLUT1 in spleens from *Pc*-infected C57BL/6 mice were also increased at 8 dpi (Figure 3G). We assessed GLUT1 expression in CD4^+^ and CD8^+^ T lymphocytes, B lymphocytes and CD11b^+^F4/80^+^/CD11c^+^/Ly6g^-^ myeloid cells by flow cytometry (gated as shown in Supplemental 1A and 1B). We found that the CD11b^+^ subset and, to a lesser extent, CD4^+^ T cells, but not CD8^+^ T and B lymphocytes, displayed increased expression of GLUT1 in response to infection (Figures 3G, 3H and Supplemental 1C). Importantly, the magnitude of GLUT1 expression in monocytic cells was ∼410%, whereas in CD4^+^ T cells was ∼29% (Figure 3H). In accordance with the expression of GLUT1 in those cell populations, the metabolic profiles of T and B lymphocytes were not altered (Figure 3I). In contrast, splenic CD11b^+^ cells displayed enhanced glycolysis in infected mice as compared to naive mice (Figure 3J).

We have previously demonstrated that infection with *P. berghei ANKA* induces a marked increase in monocyte-derived dendritic cells (MO-DCs) in the spleens of mice (*41*, *42*). We then analyzed splenic CD11b^+^F4/80^+^ cells (which are ≥90% CD11c^+^Ly6G^-^) (Supplemental 1), and selected monocytes (G1: DC-SIGN^-^MHCII^-^), inflammatory monocytes (G2: DC-SIGN^+^MHCII^-^) and MO-DCs (G3: DC-SIGN^+^MHCII^+^). As shown in Figures 3K and 3L, *Pc* infection resulted in increased frequencies of MO-DCs, and reduced percentages of inflammatory monocytes and monocytes in spleens. At 8 dpi, expression of GLUT-1 was increased in inflammatory monocytes and MO-DCs but not in monocytes (Figure 3 M). Moreover, when we compared cells from infected mice, the levels of GLUT1 were similar between MO-DCs and inflammatory monocytes, but significantly higher in MO-DCs compared to monocytes (Figure 3M). Altogether, our results suggest that activated monocytes are the primary cells responsible for increased glucose uptake in the spleens in *Pc*-infected mice.

### TNF signaling induces glycolysis in tissues from *P. chabaudi*-infected mice

As shown above, TNF signaling promotes glucose uptake in livers and spleens from *Pc*-infected mice (Figure 2K). We next asked whether TNF modulates glucose metabolism in cells from these same organs. We observed that, except for Hexokinase-3, the expression of mRNA from glycolytic enzymes (Hexokinase-1, PFKP and PKM) was increased in C57BL/6 but not TNFR^-/-^ 8-dpi (Figure 4B-E), whereas the signature of gluconeogenesis enzymes (G6PC, G6PC3, FBP1, PCK1 and PCK2) mRNA remained unchanged (Figures 4F-J). The expression of GLUT2, the main glucose transporter of hepatocytes, was similar in naïve and infected mice, both in C57BL/6 and TNFR^-/-^ mice (Figure 4K). The expression of GLUT1 protein mirrored the mRNA expression of glycolytic enzymes (Figure 4K and L), and the increased expression of GLUT1 was also lower in splenic CD11b^+^ cells from *Pc*-infected TNFR^-/-^ mice (Figure 4L). To directly assess the role of TNF during infection in this phenotype, splenocytes were stimulated with TNF *in vitro*, which reproduced the effects observed during infection, supporting a TNF-dependent regulation of GLUT1 expression (Figure 4M). In agreement with the reduced levels of GLUT1, splenic CD11b^+^ cells from TNFR^-/-^ infected mice displayed reduced ECAR when compared with CD11b^+^ from infected wild-type animals (Figure 4N), denoted as lower basal and compensatory glycolysis (Figure 4O).

**Fig. 4.**
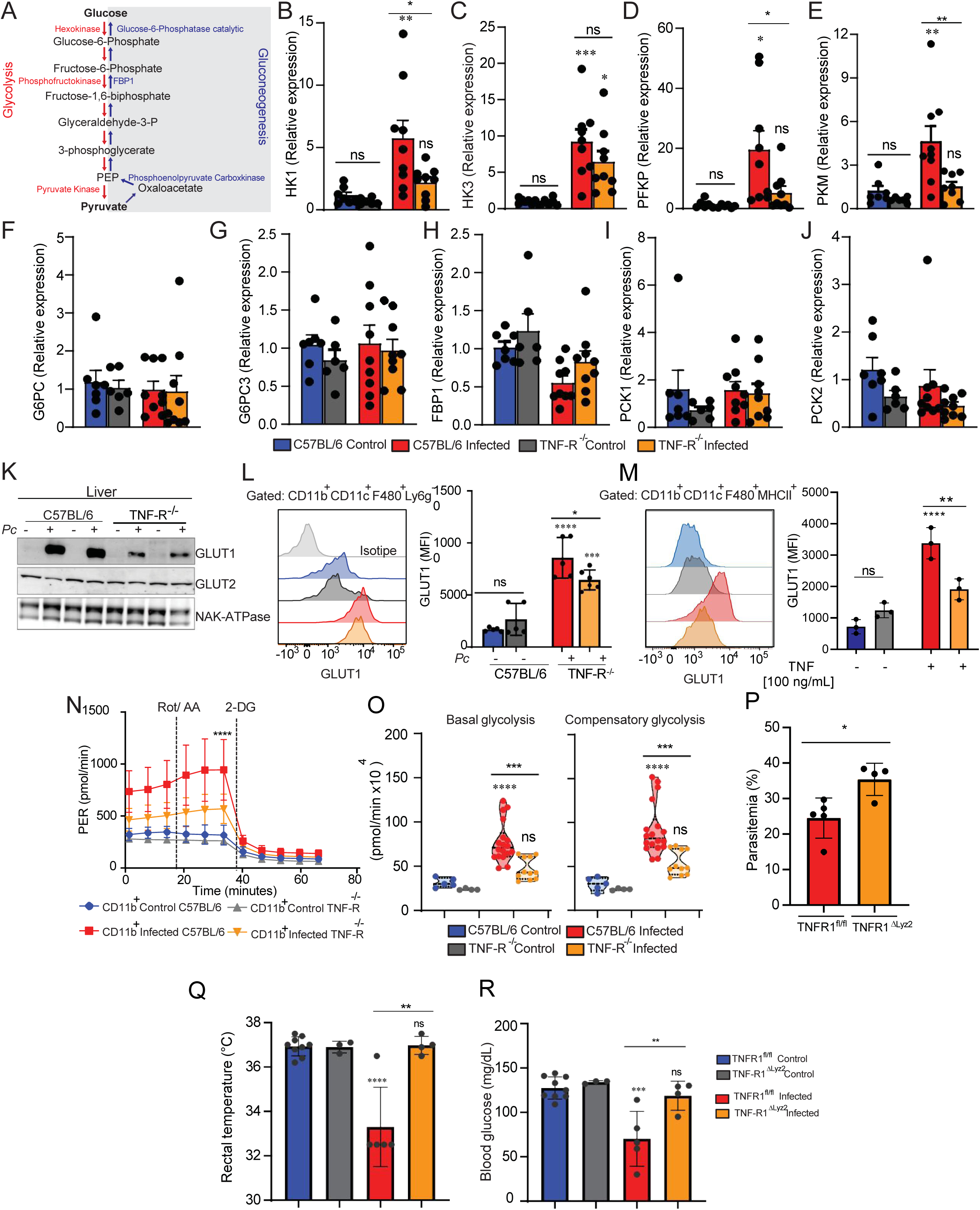
*P. chabaudi* infection triggers increased glycolysis in monocytic cells in a TNF-receptor dependent way. C57BL/6 or TNFR-/- mice were infected or not *i.p.* with *P. chabaudi* (10^5^ iRBCs). All experimental groups survived *Pc* infection throughout the observation period. (**A**) Glycolytic and gluconeogenesis pathways. Relative expression of glycolytic **(B-E)** or gluconeogenesis **(F-J)** enzymes in livers of control or infected (8 dpi) mice. **(K)** GLUT1, GLUT2 and NAK-ATPase expression in livers of infected (8 dpi) or uninfected mice evaluated by western blot. **(L)** Representative histogram and graph showing GLUT1 expression evaluated by flow cytometry in CD11b+/F4/80+/CD11c+/Ly6G-cells from spleens of infected (8 dpi) or uninfected C57BL/6 or TNFR^-/-^ mice. **(M)** Representative histogram and graph showing GLUT1 expression evaluated by flow cytometry in CD11b+/F4/80+/CD11c+/MHCII+ cells from spleens of uninfected C57BL/6 or TNFR^-/-^ mice stimulated or not with TNF (100 ng/ml) for 18h. **(N and O)** CD11b+ cells were purified from spleens harvested from infected (8 dpi) C57BL/6 (red) and TNFR-/- (orange) mice or uninfected C57BL/6 (blue) and TNFR-/- (gray) mice and cultured *ex vivo* for evaluation of ECAR by Seahorse XFe96 Analyzers. **(P)** Parasitemia was determined at 8 days-post-infection (dpi) from *TNF-R1a*ΔLyz2 and Wild-type mice. **(Q)** Rectal temperature was determined at 8 dpi in WT and *TNF-R1a*ΔLyz2 at 8 dpi. **(R)** Blood glucose levels were measured with a glucometer at 8dpi. B, C, D, E, F, G, H, I, J and M: Graphs show mean ± SEM of combined data from 2 independent experiments with 3-5 mice per group; Statistical analysis: One-way ANOVA followed by Tukey post-test. K: representative dot plot of 1 representative experiment representative of 3 independent ones; L, P, Q, R: Graphs show mean ± SEM of 1 representative experiment of at least 3 independent ones performed with 4-5 mice per group, Statistical analysis: Student’s t test. N: Graphs show mean ± SEM of 1 representative experiment of at least 3 independent ones performed with 4-5 mice per group; O: Violin plots of combined data from 3 independent experiments with 3-5 mice per group – individual dots represent the average of triplicates in each experiment – the difference refers to the number of cells isolated from naïve vs infected animals, which varied. N and O: Statistical analysis: Two-way ANOVA followed by Sidak’s post-test employing mixed effect analysis provided by Agilent Seahorse XFe96 analyzer software. Asterisks above the bars indicate comparisons between mice of the same strain before infection and at 8 dpi. Asterisks connected by horizontal lines indicate comparisons between infected groups. *P≤0.05, **P≤0.01, ***P≤0.001, **** P≤0.0001, ns = non-significant.

To further confirm that the observed effects were due to TNF signaling in myeloid cells we generated mice with conditional deletion of TNF receptor 1 in lysozyme M-expressing cells (*TNFR1*^ΔLyz2^). As expected, we observed a higher parasitemia in *TNFR1*^ΔLyz2^ mice than in WT animals (Figure 4P). Moreover, *TNFR1*^ΔLyz2^ infected mice did not decreased the rectal temperature (Figure 4Q) and blood glucose levels (Figure 4R). These data demonstrate that during experimental malaria, TNF signaling plays a key role in inducing GLUT1 expression and glucose metabolism in myeloid cells.

### TNF and HIF-1α signaling crosstalk regulates glycolytic metabolism in myeloid cells from *P. chabaudi*-infected mice

The hypoxia-inducible factor 1 alpha (HIF-1α) is an oxygen-regulated transcriptional activator that plays essential roles in mammalian development, metabolism and pathogenesis of several diseases (*43*). Its functions are primarily associated with the reprogramming of cellular energetic metabolism in response to hypoxic conditions, but in innate immune cells, HIF-1α can also promote glycolysis and quickly generate ATP and intermediate metabolites that fuel the activation and release of pro-inflammatory cytokines (*43*). Because of that, we next asked if HIF-1α is involved in the TNF-induced reprogramming of glucose metabolism in immune cells during *Pc* infection. We observed a higher nuclear expression of HIF-1α in the spleen and liver of infected mice; however, the levels of HIF-1α were lower in livers of TNFR^-/-^ compared to C57BL/6 infected mice (Figure 5A-B).

**Fig. 5.**
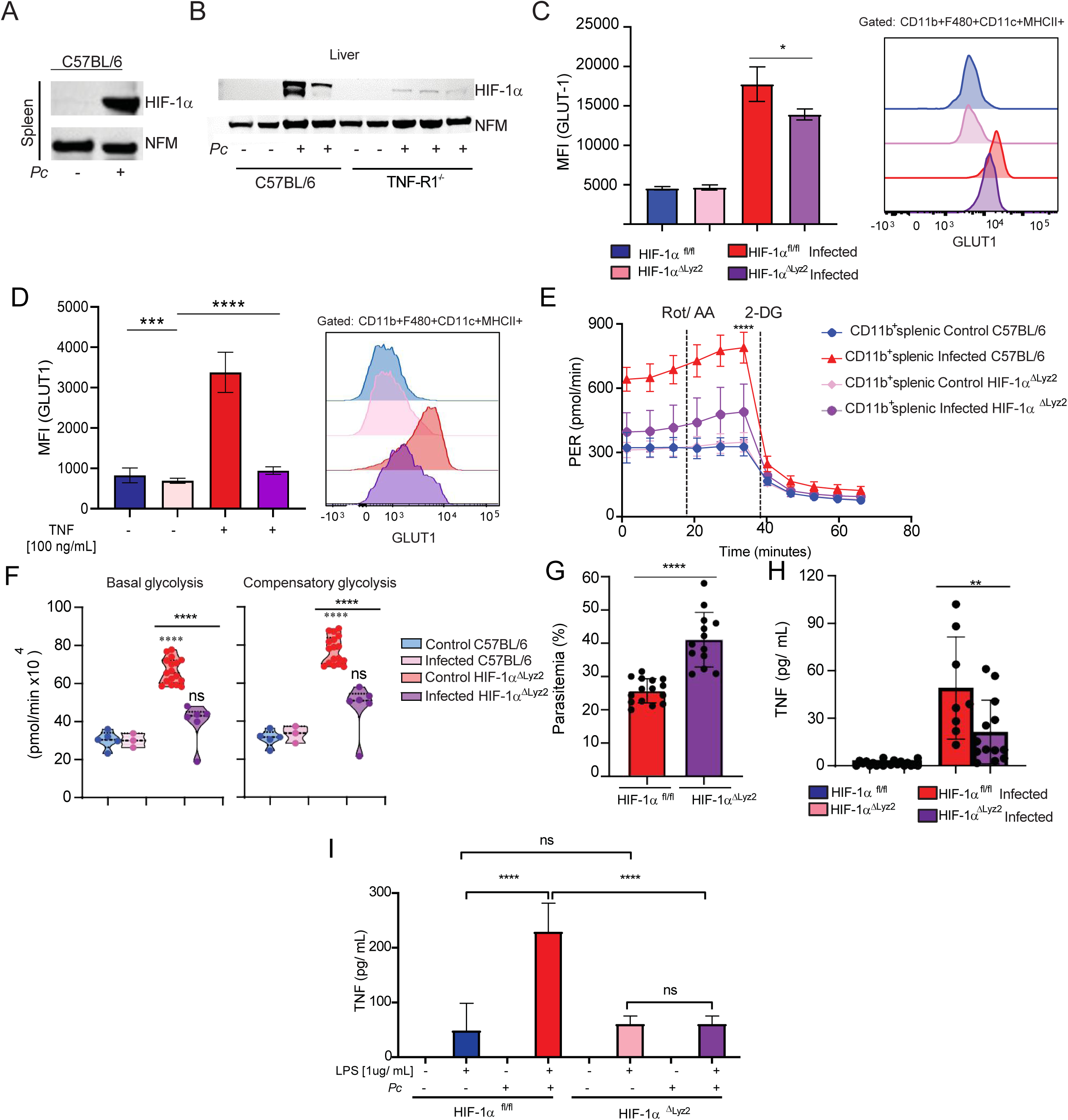
HIF-1α contributes to glycolysis induction during *Pc* infection. HIF-1α expression in the nuclear extract of cells from spleen **(A)** and liver **(B)** of infected (8 dpi) or uninfected C57BL/6 and TNFR-/- mice evaluated by western blot. (expression of nucleofosmin -NFM- was used as control). **(C)** Representative histogram and and graph showing GLUT1 expression evaluated by flow cytometry in CD11b+/F4/80+/CD11c+/MHCII+ cells from spleens of infected (8 dpi) or uninfected HIF-1aΔLyz2 and Wild-type mice. **(D)** Representative histogram and and graph showing GLUT1 expression evaluated by flow cytometry in CD11b+/F4/80+/CD11c+/MHCII+ cells from spleens of uninfected HIF-1aΔLyz2 and wild-type mice stimulated or not with TNF (100 ng/ml) for 18h. **(E and F)** CD11b+ cells were purified from spleens harvested from infected (8 dpi) WT (red) and HIF-1aΔLyz2 (pink) mice or uninfected WT (blue) and HIF-1aΔLyz2 (purple) mice and cultured ex vivo for evaluation of ECAR by Seahorse XFe96 Analyzers. **(G)** Parasitemia was determined at 8 days-post-infection (dpi) from HIF-1aΔLyz2 and Wildtype mice. **(H)** Blood TNF Levels were measured by ELISA at 8 dpi. **(I)** TNF levels in supernatants from splenic cells harvested from infected (8 dpi) or uninfected HIF-1aΔLyz2 and Wild-type mice mice and stimulated ex vivo with LPS [1ug/mL] or not during 24H. A and B: representative dot plot of 1 representative experiment representative of 3 independent ones. C: Graphs show mean ± SEM of 1 representative experiment of 3 independent ones with 3-5 mice per group; Statistical analysis: Student’s t test. D: Graphs show mean ± SEM of 1 representative experiment of 2 independent ones with 3-5 mice per group; Statistical analysis: Student’s t test. E: Graphs show mean ± SEM of 1 representative experiment of at least 3 independent ones performed with 4-5 mice per group. F: Violin plots of combined data from 3 independent experiments with 3-5 mice per group – individual dots represent the average of triplicates in each experiment – the difference refers to the number of cells isolated from naïve vs infected animals, which varied. E and F: Statistical analysis: Two-way ANOVA followed by Sidak’s post-hoc test employing mixed effect analysis provided by Agilent Seahorse XFe96 analyzer software. G and H: Graphs show mean ± SEM of combined data from 3 independent experiments with 3-5 mice per group; Statistical analysis: Student’s t test. I: Graphs show mean ± SEM of 1 representative experiment of 3 independent ones with 3-5 mice per group; Statistical analysis: One-way ANOVA followed by Tukey post-hoc test. Asterisks above the bars indicate comparisons between mice of the same strain before infection and at 8 dpi. Asterisks connected by horizontal lines indicate comparisons between infected groups. *P≤0.05, **P≤0.01, ***P≤0.001, **** P≤0.0001, ns = non-significant.

We next assessed the importance of the HIF-1α pathway for host resistance to malaria by using mice with conditional deletion of HIF-1α in myeloid cells (*HIF-1a*^ΔLyz2^) and their WT counterparts (*HIF-1a*^fl/fl^). It has been previously described that HIF-1α activity can induce the expression of GLUT1 in different mammalian cell populations (*20*). We found that glycolysis (ECAR) and GLUT1 expression were impaired, though partially, in monocytic and splenic CD11b+ cells from infected HIF-1aΔLyz2 mice (Figures 5C-5E) compared to infected WT mice. The level of GLUT1 expression that is still maintained is likely due to other host or parasite factors, such as IFN-γ (*44*). Consistent with these findings, stimulation of splenic cells with TNF *in vitro* induced a similar pattern, further supporting the link between TNF signaling and HIF-1α activation (Figure 5D). Consistent with the impaired GLUT1 expression, CD11b+ splenocytes isolated from HIF-1αΔLyz2 infected mice exhibited a marked decrease in ECAR compared to CD11b+ cells from infected control animals (Figures 5E and F). Importantly, we observed a higher parasitemia in *HIF-1a*^ΔLyz2^ mice than in WT animals (Figure 5G and Supplemental 2). At 8dpi with *Pc*, the parasitemia was higher and the circulating levels of TNF lower in *HIF-1a*^ΔLyz2^ mice compared to WT animals (Figures 5G and 5H). In addition, *Pc*-infected *HIF-1a*^fl/fl^ mice splenocytes displayed enhanced TNF secretion following LPS stimulation compared to cells from *HIF-1a*^ΔLyz2^ infected animals (Figure 5I). Altogether, these data demonstrate that the induction of the HIF-1a in myeloid cells is important for inducing glycolysis, TNF production and control of *Pc* infection.

### iNOS expression promotes HIF-1α stability, glycolytic metabolism and modulates signs of disease in *P. chabaudi*-infected mice

Our next step was to understand how TNF promotes HIF-1α expression. It is known that RNI induces HIF-1α expression in particular by enhancing the stability of this transcription factor through a mechanism that inhibits its degradation (*45*). TNF is also widely described as a major inducer of iNOS expression and RNI release by myeloid cells, such as monocytes (*46*). As expected, we observed that splenocytes from *Pc*-infected TNFR^-/-^ mice stimulated *ex vivo* with LPS released lower levels of RNI than LPS-stimulated cells from C57BL/6-infected mice (Figure 6A). Importantly, iNOS-deficient mice presented higher parasitemia than C57BL/6 animals at 8dpi with *Pc* (Figure 6B), as previously described (*47*). Similar to what was observed in TNFR^-/-^ animals, infected iNOS^-/-^ mice did show lower temperature at 8dpi with *Pc* (Figure 6C). Additionally, at this same time point, we did not observe hypoglycemia in the iNOS^-/-^ mice infected with *Pc* (Figure 6D). As expected, splenic CD11b^+^ cells from iNOS^-/-^ infected mice displayed higher OCR when compared to C57BL/6 infected with *Pc* (Figure 6E), indicating increased basal and maximal mitochondrial respiratory capacities in the absence of iNOS (Figure 6F).

**Fig. 6.**
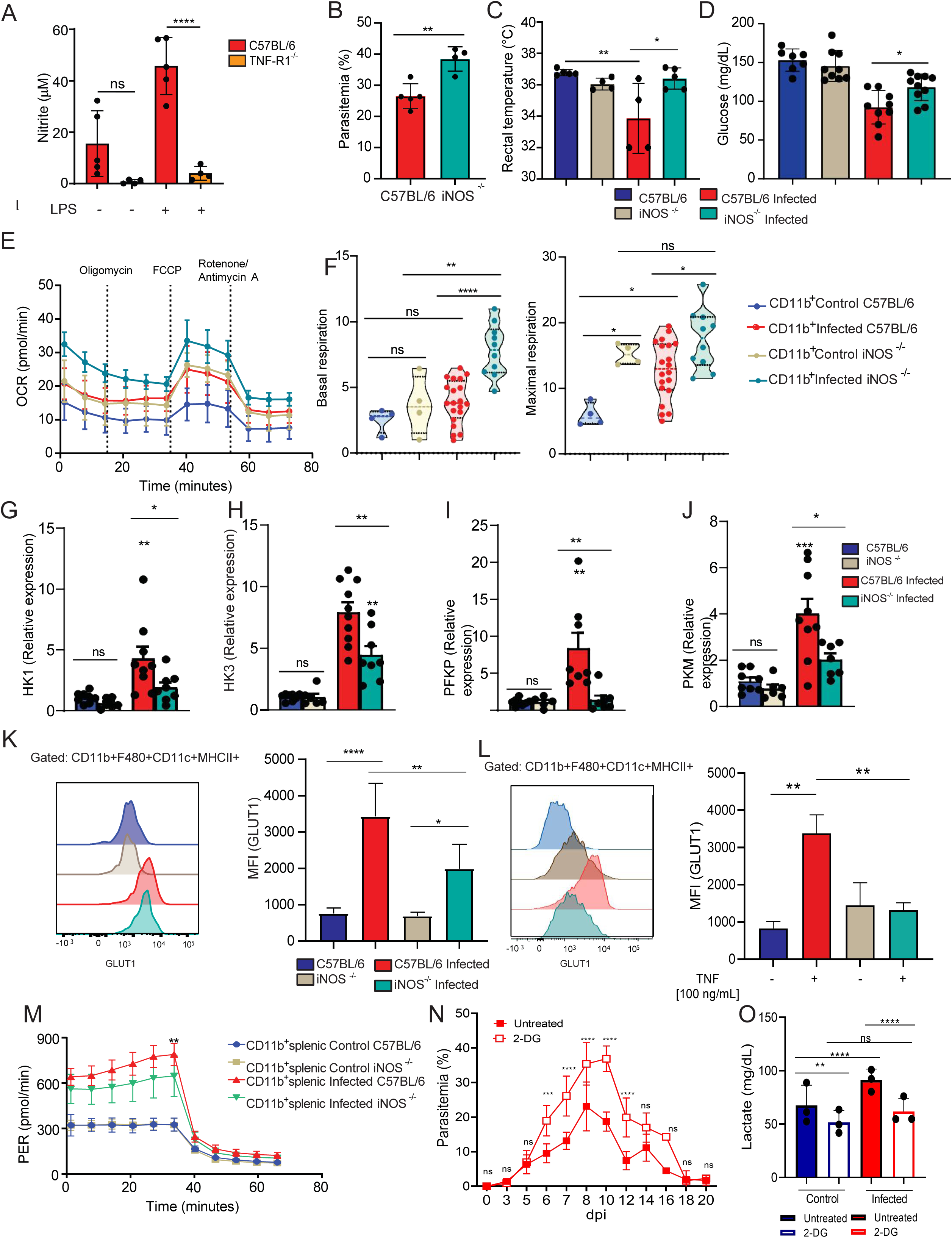
Malaria disease parameters in iNOS deficient mice mirror those observed in TNF receptor deficient animals. **(A)** Nitrite levels in supernatants from CD11b+ splenic cells harvested from infected (8 dpi) C57BL/6 or TNF-R1-/-mice and stimulated ex vivo with LPS [1ug/mL] or not during 48H. **(B)** Parasitemia was determined at 8 days-post-infection (dpi) from C57BL/6 and iNOS knockout mice. **(C)** Rectal temperature was determined at 8 dpi. **(D)** Blood glucose levels were measured with a glucometer at 8dpi. **(E and F)** CD11b+ cells were purified from spleens harvested from infected (8 dpi) C57BL/6 (red) and iNOS knockout (green) mice or uninfected C57BL/6 (blue) and iNOS knockout (beige) mice and cultured ex vivo for evaluation of ECAR by Seahorse XFe96 Analyzers. **(G-J)** Relative expression of glycolytic enzymes in livers of naïve or infected (8 dpi) mice. **(K)** Representative histogram and graph showing GLUT1 expression evaluated by flow cytometry in CD11b+/F4/80+/CD11c+/MHCII+ cells from spleens of infected (8 dpi) or uninfected C57BL/6 or iNOS^-/-^ mice. **(L)** Representative histogram and graph showing GLUT1 expression evaluated by flow cytometry in CD11b+/F4/80+/CD11c+/MHCII+ cells from spleens of uninfected C57BL/6 or iNOS^-/-^ mice stimulated or not with TNF (100 ng/ml) for 18h. **(M)** CD11b+ splenic cells were purified from infected C57BL/6 (red - 8 dpi), iNOS^-/-^ (green - 8 dpi), and uninfected mice and cultured ex vivo for quantification of proton efflux rate (PER) using glycolytic rate assay by Seahorse XFe96. **(N)** Parasitemia was determined during 20 days-post-infection (dpi) from wild-type infected mice treated with 2-[14C]-deoxyglucose (250mg/kg daily from 4 to 15 dpi) or PBS. **(O)** Lactate was measured in the plasma of mice. A: Graphs show mean ± SEM of 1 representative experiment of 3 independent ones with 4-5 mice per group; Statistical analysis: One-way ANOVA followed by Tukey post-hoc test. B, C and D: Graphs show mean ± SEM of 1 representative experiment of 3 independent ones with 3-5 mice per group (B and C) or of combined data from 2 independent experiments with 3-5 mice per group (D); Statistical Analysis: Student’s t test. E: Graphs show mean ± SEM of 1 representative experiment of at least 3 independent ones performed with 4-5 mice per group. F: Violin plots of combined data from 3 independent experiments with 3-5 mice per group – individual dots represent the average of triplicates in each experiment – the difference refers to the number of cells isolated from naïve vs infected animals, which varied. E and F: Statistical analysis: Two-way ANOVA followed by Sidak’s post-hoc test employing mixed effect analysis provided by Agilent Seahorse XFe96 analyzer software. G, H, I, J, N, O: Graphs show mean ± SEM of combined data from 2 independent experiments with 3-5 mice per group; Statistical Analysis: Student’s t test: One-way ANOVA followed by Tukey post-hoc test. K: Graphs show mean ± SEM of 1 representative experiment of 3 independent ones with 3-5 mice per group; Statistical analysis: Student’s t test. L: Graphs show mean ± SEM of 1 representative experiment of 2 independent ones with 3-5 mice per group; Statistical analysis: Student’s t test. M: Graphs show mean ± SEM of 1 representative experiment of at least 3 independent ones performed with 4-5 mice per group; Statistical analysis: Two-way ANOVA followed by Sidak’s post-hoc test employing mixed effect analysis provided by Agilent Seahorse XFe96 analyzer software. Asterisks above the bars indicate comparisons between mice of the same strain before infection and at 8 dpi. Asterisks connected by horizontal lines indicate comparisons between infected groups. *P≤0.05, **P≤0.01, ***P≤0.001, **** P≤0.0001, ns = non-significant.

In accordance with the results obtained in *Pc-*infected TNFR^-/-^ mice, iNOS deficiency resulted in reduced hepatic expression of the glycolytic enzymes HK1, HK3, PFKP and PKM (Figures 6 G-J) as well as reduced GLUT1 expression in splenic monocytes (Figure 6K). TNF stimulation *in vitro* resulted in a similar profile to that observed *in vivo*, further supporting a TNF–iNOS–dependent regulation of glycolytic pathways (Figure 6L). Accordingly, glycolytic metabolism in splenic monocytic cells of iNOS^-/-^ infected animals was decreased compared to those of C57BL/6 infected mice, as denoted by a lower ECAR (Figure 6M). Finally, to directly link glycolytic metabolism and TNF production during *Pc* infection, we inhibited glycolysis *in vivo* using 2-deoxy-D-glucose (2-DG). Treatment with 2-DG resulted in a marked increase in parasitemia compared to untreated infected mice (Figure 6O), resembling the impaired parasite control observed in HIF-1αΔLyz2, TNFR^-/-^ and iNOS^-/-^ mice. Consistent with effective glycolytic blockade, 2-DG–treated mice displayed reduced lactate levels during infection (Figure 6N and O). Thus, the inhibition of glycolysis mirrored the metabolic and immunological alterations observed upon disruption of the TNF–HIF-1α–iNOS axis, supporting the conclusion that this pathway is critical for sustaining glycolytic metabolism, TNF production and effective parasite control during *Pc* infection. Altogether, these data demonstrated that RNI induces HIF-1α expression, glycolysis, and TNF release by monocytic cells, leading to control of parasitemia but also promoting clinical signs of disease, such as hypothermia and hypoglycemia, in *Pc*-infected mice.

## Discussion

Malaria disease manifestations include severe metabolic changes in the host organism. Different studies report that decreased glycaemia and increased lactate plasma levels followed by acidosis are common manifestations in patients with severe malaria (*5*, *48–50*). However, the mechanisms that lead to hypoglycemia and increased lactate levels during malaria are poorly understood. A better understanding of the mechanisms that mediate host metabolic alterations and the connections of such events with the immune response against the parasite may provide new insights for therapeutic interventions in malaria patients. In this study, we found that TNF signaling mediates changes in host energy metabolism, accompanied by increased expression of GLUT1 and enhanced glycolysis in monocytes. In addition, we found that TNF-induced RNI stabilizes the transcription factor HIF-1α, which promotes both a metabolic shift towards glycolysis and the expression of pro-inflammatory cytokines by monocytes. Furthermore, we report that TNF, iNOS and HIF-1α have an important role in controlling *Pc* infection and signs of systemic inflammation in this mouse malaria model.

Previous studies have shown that TNF induces the production of RNI through the upregulation of iNOS via the NF-κB pathway (*51–54*). TNF-mediated iNOS expression is critical for RNI production, which in turn stabilizes HIF-1α by inhibiting prolyl hydroxylases (PHDs) even under normoxic conditions (*52*). HIF-1α then upregulates the expression of glycolytic genes, including GLUT1 (*55–57*). TNF has been described as a critical mediator in malaria, driving cytokine release and parasitemia control (*58*). It also enhances glucose uptake in tissues, aligning with our findings of increased glycolysis in monocytes (*59*). The role of iNOS in malaria is well documented. IFN-γ and TNF induced RNI inhibits parasite growth, but can cause tissue damage and organ dysfunction, especially in severe malaria (*60*). Recent studies also highlight the complexity of glycemia regulation during *Plasmodium* infection describing its role in modulating parasite virulence and transmission (*61*). These studies demonstrate the critical function of TNF and iNOS in immune responses against *Plasmodium*. Aligning with our findings that this axis and metabolic rewiring are essential for monocyte activation and outcome of *Pc* infection

Interestingly, we found that glucose uptake was significantly increased in spleens and livers of *Pc*-infected mice and that this effect was partially dependent on TNF signaling. Consistently, TNF promoted increased expression of GLUT1 and glycolytic energy metabolism in non-parenchymal monocytic cells, but not hepatocytes or lymphocytes in livers and spleens, respectively. These results indicate that the hypoglycemia of *Pc*-infected mice is, at least in part, caused by TNF-driven GLUT1-mediated enhanced glucose uptake and glycolysis in monocytes. Insulin is the major hormone responsible for the upregulation of glucose uptake in several organs, including adipose and skeletal muscle tissues, which happens mainly through the upregulation of GLUT4 (*62*, *63*). In mice, GLUT1 expression is recognized as independent of insulin, in contrast to GLUT4 (*64*). In our model, this regulation appears to be driven by pro-inflammatory cytokines, particularly TNF. Supporting this, our results show that *in vitro* stimulation with TNF, significantly increases GLUT1 expression in monocytes, accordingly to the *ex vivo* phenotype observed in infected animals. Altogether, our findings suggest that the increase in glucose uptake and glycolysis in monocytes is insulin-independent. Although the frequency of MO-DCs increases during infection, other cell populations may also contribute to glucose consumption. Further experiments, including the assessment of GLUT1 function in these populations, are needed to clarify their relative contribution to glucose consumption during infection

Following the rupture of parasitized red blood cells, there is a release of pathogen and danger-associated molecular patterns that activate Toll-like receptors (TLRs), Nod-like receptors (NLRs), and the cyclic GMP–AMP (cGAS) synthase (*10*, *65*). Activation of these innate immune receptors and sensors triggers the production of high levels of pro-inflammatory cytokines, such as TNF, IL-6, IL1β and IL-12, all of which participate in host immunity against *Plasmodium* infection (*66*). The importance of the glycolytic pathway for innate immune cell activation has been demonstrated in different studies (*20*, *67*). For instance, monocyte differentiation from a resting to an inflammatory state requires a shift in energy metabolism to high glucose consumption and rapid energy generation by glycolysis (*68*). Activated monocytes, along with dendritic cells, then produce IL-12, which is critical for the differentiation of Th1 lymphocytes that produce IFN-γ, a cytokine that activates splenic macrophages to eliminate *Plasmodium*-infected red blood cells (*10*, *66*).

Importantly, we found that similarly to TNFR^-/-^ mice, iNOS-deficient mice display impaired glycolytic metabolism in monocytic cells from *Pc*-infected mice. RNI are produced in macrophages once they receive two signals. The first signal is the priming with IFN-γ. The second signal is the activation of monocytes with TNF induced by microbial products via TLRs. Like in TNFR^-/-^ mice, iNOS deficiency resulted in increased parasitemia following *Pc* infection (*69*, *70*). HIF-1α is required for the optimal expression of many glycolytic genes in macrophages, including those encoding GLUT1, hexokinase, phosphofructokinase and pyruvate kinase (*43*). Recently, it was suggested that RNI promotes *S*-nitrosylation at the Cys533 residue of HIF-1α molecules (*71*), promoting HIF-1α stabilization by direct inactivation of PHDs (*72*). Indeed, the impairment of RNI release during *Pc* infection, resulted in reduced translocation of HIF-1α to the nuclei of splenic monocytes and non-parenchymal liver cells, which was markedly increased in infected WT mice. Therefore, we propose that the mechanism by which TNF signaling regulates HIF-1α stabilization/expression during *Pc* infection is mediated by RNI.

Different studies have demonstrated that HIF-1α is critical for host resistance to different pathogens (*73*), but both the participation of this transcription factor in malaria as well as the role of glycolytic metabolism in resistance to *Plasmodium* infection have not been explored. The *HIF-1α*^ΔLyz2^ animals, which are genetically deficient for HIF-1α and have impaired glycolysis only in myeloid cells, were more susceptible to *Pc* infection, further demonstrating the importance of myeloid cells in controlling parasitemia. This might be explained by the importance of glycolysis for the release of TNF and other pro-inflammatory cytokines by monocytes that are activated during *Plasmodium* infection (*3*, *40*). Although we have found that *Pc* infection-induced increase in nuclear translocation of HIF-1α was impaired in the absence of TNF signaling, *HIF-1α*^ΔLyz2^ animals also displayed lower TNF secretion in response to *Pc* infection than WT mice. Therefore, these results indicate that the in vivo relationship between TNF signaling and HIF-1α-driven glycolysis is complex and that a positive feedback loop between TNF production, HIF-1α expression and glycolysis must exist during *Pc* infection.

The link between TNF and HIF has been explored in different cell types. Previously, studies demonstrated a crosstalk between NF-kB and HIF-1α. TNF signaling recruits different intracellular adaptors that activate multiple signal transduction pathways. One consists of the TRAF family of proteins, which can lead to IKK-dependent activation of NF-kB, thereby promoting inflammatory responses (*74*). Conversely, HIF-1α can also regulate NF-kB activation. Koedderitzch and collaborators (*75*) have recently shown that TNF induces glycolytic shift via GLUT-1 and HIF-1α in rheumatoid arthritis (RA). As with malaria, TNF is a pivotal cytokine involved in the pathogenesis of RA. *In vitro* treatment of fibroblast-like synoviocytes with TNF induces upregulation of GLUT-1 and HIF-1α, which depends on TAK1-induced NF-kB activation downstream of TNF receptor 1 (*75*). In addition, NF-kB plays a central role in inflammatory responses, orchestrating the expression of TNF, IL-6, adhesion molecules and enzymes. Moreover, an NF-kB binding site is present in the proximal promoter site of the HIF-1α gene, indicating that NF-kB activation can regulate its expression (*76*, *77*).

*Plasmodium* parasites do not synthesize glucose, being solely dependent on host glucose as a source of energy required for parasite replication. On the one hand, the increased glucose uptake and subsequent glycolytic metabolism in monocytes promote hypoglycemia, helping to starve the erythrocytic stage of the parasite, controlling its replication. On the other hand, hypoglycemia may also be detrimental to the host. For instance, lactate, the final product of glycolytic metabolism of glucose, when secreted in high amounts, causes acidosis with decreased blood pH and severe consequences to the host, such as extreme fatigue, pain, overall feelings of physical discomfort and decreased appetite (*5*).

Indeed, we found that the absence of TNF signaling and iNOS^-/-^ mice reverted the development of hypothermia, which is a known sickness manifestation in *Pc* infection-induced experimental malaria in mice (*78–80*). Moreover, the attenuation of clinical symptoms in TNFR^-/-^ infected mice extended beyond thermal regulation. Despite increased parasitemia, these animals displayed restored physical activity, food consumption, energy expenditure and respiratory exchange rate compared to WT animals. Although restored physical activity, food consumption and energy expenditure in knockout mice may contribute to the observed systemic metabolic parameters by altering energy balance, these effects are not mutually exclusive with the TNF-driven, cell-intrinsic metabolic mechanisms described here. These findings demonstrate that TNFR^-/-^ mice are more “tolerant” to disease. Hence, our results allow us to conclude that although TNF promotes host resistance during malaria, it plays a detrimental role during *Plasmodium* infection.

In summary, our results indicate that TNF-induced RNI induces HIF-1α-mediated glycolysis and pro-inflammatory response by monocytes, promoting host resistance to infection with *Plasmodium*. However, this exacerbated activation of monocytes may be detrimental to the host. Hence, our findings enlighten a fundamental mechanistic question related to the pathophysiological role of *Plasmodium* infection in the vertebrate host. These findings may provide insights for developing novel therapeutic interventions to treat this devastating disease.

## Materials and Methods

### Ethics statement

All experiments were carried out in accordance with institutional guidelines for animal ethics and approved by the Ethics Committee on Animal Use of Ribeirão Preto Medical School University of São Paulo (CEUA 94 / 2020), Institutional Ethics Committees from Oswaldo Cruz Foundation (Fiocruz-Minas, CEUA/LW15/14, and LW16/18) and UMMS (IACUC A-1369-14-5), respectively.

### Mice

C57BL/6 and TNFR^-/-^ (TNFR p55/p75 chains double knockout mice) were obtained from the Center for Breeding of Transgenic Mice from the Ribeirao Preto Medical School. Conditional knockout mice (*TNFR1*^ΔLyz2^) were generated by crossing LysMCre^+/-^ to *TNFR1*^flfl^ mice, which were maintained on a C57BL/6 genetic background. Mice used in experiments were sex and age-matched: female mice between 8–12 weeks of age. All mouse lineages were housed in micro-isolators in a maximum number of six mice per cage, in a specific pathogen-free facility at the Oswaldo Cruz Foundation, or Ribeirão Preto Medical School or University of Massachusetts Medical School, under controlled temperature (22–25°C) and 12-h light-dark cycle and provided with water and food ad libitum.

### Experimental Infections

*Plasmodium chabaudi chabaudi* AS strain (*Pc*) was used for experimental infections (*81*, *82*). First, *Pc* was maintained in C57BL/6 mice by serial passages once a week up to eight times. For experimental infection, mice were injected intraperitoneally (i.p.) with 10^5^ infected red blood cells (iRBCs) diluted in 100 µl PBS-1X. The percentage of RBCs containing parasites (parasitemia) was measured in the blood with Giemsa at different days post-infection (dpi). Infected and non-infected mice were treated daily with a glycolytic inhibitor 2-deoxy-D-glucose (2-DG) at a dose of 250 mg/kg, administered from day 4 to day 15 post-infection (dpi). Control groups, including both infected and non-infected animals, received PBS 1X following the same treatment schedule.

### Glucose Measurement

The glucose uptake was measured using 2-[14C]-deoxyglucose. Experiments were performed at the National Mouse Metabolic Phenotyping Center (MMPC) at UMASS Medical School. At 6 am on day 8 dpi, basal glucose uptake in individual organs was measured using an intravenous injection of 2-[14C]-deoxyglucose. After 1h, mice were anesthetized, and tissue samples were taken for organ-specific levels of 2-[14C]-deoxyglucose-6-phosphate. Blood glucose levels were measured with an *Accu-chek* glucometer.

### Mouse RNA-Seq

RNA-seq was performed in biological replicates (3 mice per group). Liver samples were collected from C57BL/6 mice at different time points: 12 am; 6 am; 12 pm, and 6 pm from mice at day 8 post-infection with *P. ch*. or uninfected control. RNA-seq libraries were prepared using the TruSeq Stranded mRNA Kit (*Illumina*) following the manufacturer’s instructions. Briefly, poly-A-containing mRNA molecules were purified using poly-T oligo-attached magnetic beads and fragmented using divalent cations. The RNA fragments were transcribed into cDNA using SuperScript II Reverse Transcriptase (*Invitrogen*), followed by second strand cDNA synthesis using DNA Polymerase I and RNase H. Finally, cDNA fragments then have the addition of a single ‘A’ base and subsequent ligation of the adapter. The products were then purified and enriched by PCR using paired-end primers (*Illumina*) for 15 cycles to create the final cDNA library. The library quality was verified by fragmentation analysis (*Agilent Technologies 2100 Bioanalyzer*) and submitted for sequencing on the Illumina NextSeq 500 (*Bauer Core Facility Harvard University*). These samples were collected in according to the circadian cycle published by our group (*32*).

### RNA extraction and Real-time PCR

Liver samples were collected from C57BL/6 mice at 8 dpi with *P. ch.* or uninfected controls. Tissue fragments were stabilized in RNAlater (Qiagen) and RNA extraction was performed using an RNeasy kit (Qiagen) according to manufacturer instructions. A total of 1mg of RNA was converted to cDNA using iScript cDNA Synthesis Kit (Bio-Rad). Real-time PCR was performed with the SYBR Green master mix (Bio-Rad). Primer sequences are: CCCTCACACTCAGATCATCTTCT (Foward primer) and GCTACGACGTGGGCTACAG (Reverse primer) (NM_013693).

### Isolation of splenocytes and purification of CD11b^+^, B and T cells

Spleens from control and infected mice at 8 dpi were dissociated through a 100-μm nylon cell strainer for obtaining single-cell suspensions. For *ex vivo* cultures, known concentrations of splenocytes were resuspended in RPMI 1640 medium supplemented with penicillin, streptomycin, and 10% fetal bovine serum (FBS) (*Gibco, ThermoFisher*). In some experiments, splenocytes (2 × 10^6^ cells) were stimulated in vitro with 1 μg/ml of LPS for 24h, or TNF-α [20ng/ml] or not during 15 minutes. For CD11b+ cell-, B cell- or T cell-purification, immunomagnetic beads for positive selection of CD11b^+^ cells, B cells and T cells (*Miltenyi Biotec*) were used, respectively, according to manufacturer’s instructions.

### Hepatocyte’s isolation

First, the liver portal vein was cannulated with a 24G catheter. The inferior cava vein was cutted to allow perfusion flow. Perfusion was done with 50 mL of Hanks A and then 25 mL of Hanks B with 20 mg of collagenase. After that, tissue was dissociated and samples were filtered in a 40 mM membrane. Cells were centrifuged: 60x g (5 min, 4oC), disposing of supernatants, 60x g (5 min, 4oC), and resuspended in cold RPMI 10% SFB.

### Liver non-parenchymal-cells

Livers were first digested in a collagenase solution (5 mg/liver in 10 mL of RPMI 2% SFB) at 37oC for 35 min. Then, the solution was added PBS/BSA 0,5% + EDTA in order to inactivate collagenase and cells were differentially separated by centrifugation: 300 x g (10 min, 4oC) disposing supernatants, 60 x g (3 min, 4oC) twice preserving supernatants and finally, 300 x g (10 min, 4oC) disposing supernatants. The samples were filtered through 100 μm cell strainers and centrifuged for 300 x g (10 min, 4oC). Red blood cells were lysed using ACK (ammonium chloride potassium) buffer. Concentrations of single-cell suspensions were then adjusted following counting on the hematocytometer.

### Western blotting

Mouse splenocytes, magnetically purified CD11b+ splenic cells, isolated hepatocytes, isolated liver non-parenchymal cells, or total liver samples from infected or non-infected mice were lysed using RIPA buffer (Sigma) supplemented with protease and phosphatase inhibitor cocktail (Thermo Fisher Scientific).After 15 min on ice, lysates were centrifuged at 13,000 × g for 10 min at 4 °C. The proteins were separated in a 10%-acrylamide NUPAGE *Bis-tris* Protein gels (*Invitrogen*) and transferred onto nitrocellulose membranes. Membranes were blocked with 5% (wt/vol) nonfat milk (*Molico*) in Trisbuffered saline with 0.1% Tween-20 (TBST) for 1 h at room temperature and then incubated overnight at 4°C with primary antibodies. The membranes were incubated with GLUT1 (*Abcam*, 1:1000), GLUT2 (*Abcam*, 1:1000), Na,K-ATPase (*Cell Signaling, 1:2000)* and β-actin (Sigma, 1:5000) specific antibodies. Subsequently, membranes were repeatedly washed with TBST and then incubated for 2 h with the anti-Rabbit HRP-conjugated secondary antibody (1:30.000 dilution; *Sigma-Aldrich*). Immunoreactivity was detected with Clarity Max ECL Substrate (Biorad) and then the chemiluminescence signal was recorded on the iBright FL1500 (*ThermoFisher*). Data were analyzed with iBright FL1500.

### Metabolic Profiling

Metabolic profiling of CD11b^+^ cells, B cells, T cells, hepatocytes, and non-parenchymal cells was undertaken using a Seahorse XFe96 Extracellular Flux Analyzer (*Agilent*) in microplates. Basal oxygen consumption rate (OCR) and extracellular acidification production (ECAR) were measured by Phenotypic Seahorse Kit (KIT 103325-100), or Seahorse XF Cell Mito Stress Tests (KIT 103015-100), or Glycolytic Rate Assay Kit (KIT 103344-100) according to the manufacturer’s instructions.

### Flow cytometry

Splenocytes (2 × 10^6^ cells) from mice at 0 (uninfected controls) and 8 days post-infection were stained with fluorophore-labeled monoclonal antibodies (mAbs) specific for cell surface markers. The following flow cytometry specific mAbs were used: CD11c-PECy7 clone N418 (1:200), CD11b-APCCy7 clone M1/70 (1:200), Ly6c PERCP clone H.K1.4 (1:200), F4/80-PE clone BM8 (1:200), Ly6g FITC clone 1A8 (1:200), CD45 Pacific Blue-clone30F-11 (1:200), CD3 FITC clone 145-2c11 (1:200), CD4 PE clone RM4-5 (1:200), CD8 PERCP clone 53-6.7 (1:200), CD19 PeCy7 clone 6D5(1:200) and Bv510 (Live/Dead kit, ThermoFisher). Intracellular staining of GLUT1 Alexa Fluor 647 [EPR3915] (1:100) was performed following cell fixation and permeabilization using the eBioscience Foxp3 Fixation/Permeabilization Kit. Data were acquired in a FACSCanto II machine (BD Biosciences) and analyzed using FlowJo software (Tree Star).

### Cytokine measurement

Plasma was collected from whole blood and supernatants were collected from cell cultures. Cytokine concentration in these samples were determined using CBA mouse Inflammation Kit (BD^TM^) or ELISA according to the manufacturers’ instructions (*Biolegend*).

### Food Intake and Physical Activity

Mouse food intake and physical activity were assessed for 3 days prior to infection and at days 6, 7 and 8 dpi with *Pc* using metabolic cages (*TSE Systems*, Chesterfield, MO) located at the National Mouse Metabolic Phenotyping Center (MMPC) at UMASS Medical School. Mice were housed under controlled temperature and lighting, with food and water ad libitum. Food intake and physical activity monitoring was fully automated using the *TSE Systems LabMaster* platform*. LabMaster* cages allowed the use of bedding, thus, minimizing any animal anxiety during the experimental period. Physical activity was calculated based on quantitative measurement of horizontal and vertical movement (XYZ-axis). In addition, non-invasive measures of O_2_ consumption and CO_2_ production were used to calculate the respiratory exchange ratio to reflect energy expenditure.

### RNA-Seq Analysis

Gene expression of each experimental group was shown at different time points 12 am; 6 am; 12 pm and 6 pm. The average expression of all time points was calculated for visualization in heatmaps. The heatmaps presented in Figures 1 and 2 were developed with a free version of the software Morpheus (https://software.broadinstitute.org/morpheus). For hierarchical clustering, we used linkage from the average expression of each gene in metric One Minus Pearson Correlation.

### Statistical analysis

*GraphPad Prism 7.0 software* was used for statistical analysis. Multiple-group comparisons were performed with either one-way ANOVA (followed by Tukey’s post hoc test) or two-way ANOVA (followed or not by Sidak’s post hoc test). Unpaired two-tailed Student’s t-test was used for the comparison of two conditions. Results are expressed as means ± SEM. P value ≤ 0.05 was considered significant.

## Supporting information

Supplemental figures 1, 2 and 3 and its subtitle

## Acknowledgments

The authors thank members of the D.L.C., J.S.S., J.C.A.F. and R.T.G. groups for scientific discussions and technical help. Denise Brufato Ferraz and Francielle Pioto for the excellent technical assistance. Cristiane Gomes and Patricia Palhares for excellent financial technical assistance.

## Author contributions

K.C.M., J.C.A.F. and R.T.G. designed, K.C.M., N.P.S.L. P.A.A., I.C.H., L.G.V., J.E.T.K., D.L.C. performed experiments. I.C.H, P.A.A., O.O., N.P.S.L., K.C.M. and R.T.G. analyzed data, D.L.C., J.S.S., J.C.A.F. and R.T.G. contributed with reagents, materials, and/or analysis tools, and K.C.M., N.P.S.L. and R.T.G. wrote the manuscript. All authors revised and approved the final version of the manuscript.

## Data and materials availability

GEO (https://www.ncbi.nlm.nih.gov/geo/query/acc.cgi?acc=GSE109908) accession number is GEO: GSE109908.

