## Supplemental figures 1, 2 and 3 and its subtitle for "Reprogramming of host energy metabolism mediated by the TNF-iNOS-HIF-1α axis plays a key role in host resistance to *Plasmodium* infection"

***Contact info**

Gazzinelli, Ricardo T and Kely Catarine Matteucci.

**Supplemental 1**: Gating strategies for flow cytometry analysis from spleens.

(A) Lymphoid cells: B lymphocytes (Live/ Lymphocytes/ $CD45^+$/ $CD19^+$), $CD4^+$ T lymphocytes (Live/ Lymphocytes/ $CD45^+$/ $CD3^+$/ $CD4^+$) and $CD8^+$ T lymphocytes (Live/ Lymphocytes/ $CD45^+$/ $CD3^+$/ $CD8^+$). (B) Myeloid monocytic cells: (Live/ $CD45^+$/ $F4/80^+$/ $CD11b^+$ $CD11c^+$/ $Ly6G^-$). (C) Representative histogram of GLUT1 expression that was evaluated by flow cytometry in splenic cells from infected (red – 8 dpi) and uninfected (blue) mice.

**Supplemental 2** HIF1α deficiency results in higher parasitemia following *P. chabaudi* infection. Parasitemia from wild type (pink) and *HIF-1a*ΔLyz2 (purple) infected mice was determined at 0, 3, 5, 8, 10, 12, 14, 16, 18, 21 and 26 days-post-infection (dpi). ***P≤0.001 (t-test).

**Supplemental 3**: Densitometric analysis of GLUT expression and HIF-1α levels in liver and splenic cell populations. (A)

Representative immunoblot and densitometric quantification of GLUT1 protein levels in whole liver extracts from control and infected C57BL/6 mice, normalized to β-actin. (B) Immunoblot analysis and corresponding densitometry of GLUT1 expression in total liver homogenates and isolated hepatocytes from control and infected C57BL/6 mice. (C) GLUT1 protein levels in non-parenchymal liver cells from control and infected C57BL/6 mice, quantified by densitometric analysis and normalized to β-actin. (D) Immunoblot detection and densitometric quantification of GLUT1 expression in splenic T cells, B cells, CD11b$^{+}$ cells, and total spleen lysates from control and infected C57BL/6 mice. (E) Representative immunoblots and densitometric quantification of GLUT1 and GLUT2 expression in liver samples from control and infected C57BL/6 and TNFR$^{-}$ /$^{-}$ mice, normalized to Na$^{+}$ /K$^{+}$ -ATPase. (F) Immunoblot and densitometric analysis of HIF-1α protein levels in spleen lysates from control and infected C57BL/6 mice, normalized to NFM.(G) Representative immunoblot of HIF-1α protein levels in liver extracts from control and infected C57BL/6 and TNFR1$^{-}$ /$^{-}$ mice, with NFM used as a loading control. The accompanying bar graph shows densitometric quantification of HIF-1α normalized to NFM. Bars represent control and infected conditions for each genotype, and statistical comparisons are indicated as shown.Asterisks connected by horizontal lines indicate comparisons between infected groups. *$P \leq 0.05$, **$P \leq 0.01$, ***$P \leq 0.001$, **** $P \leq 0.0001$, ns = non-significant.

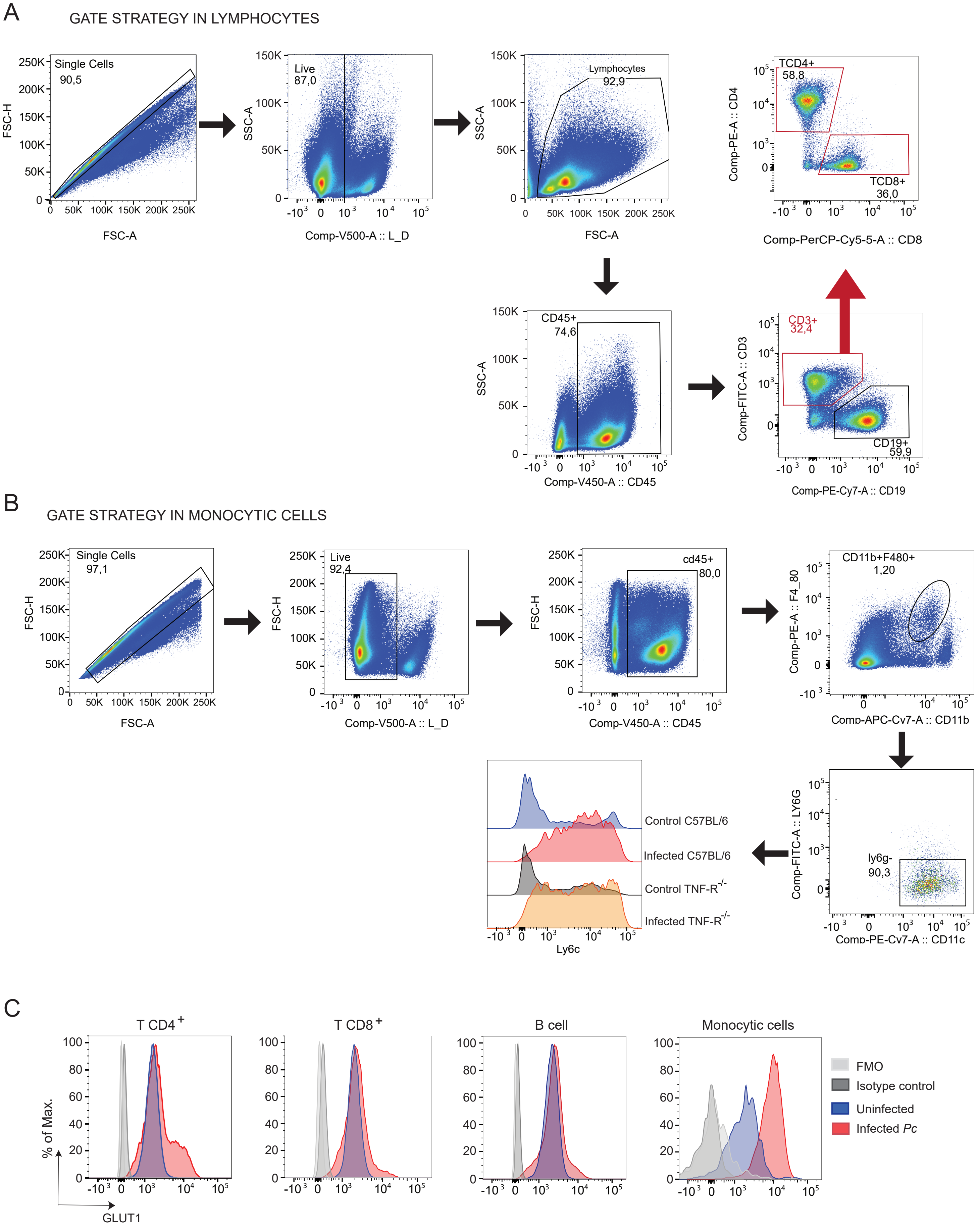

A
GATE STRATEGY IN LYMPHOCYTES
Single Cells
90,5
FSC-H
FSC-A
Live
87,0
SSC-A
Comp-V500-A :: L_D
Lymphocytes
92,9
SSC-A
FSC-A
TCD4+
58,8
TCD8+
36,0
Comp-PE-A :: CD4
Comp-PerCP-Cy5-5-A :: CD8
CD45+
74,6
SSC-A
Comp-V450-A :: CD45
CD3+
32,4
CD19+
59,9
Comp-FITC-A :: CD3
Comp-PE-Cy7-A :: CD19
B
GATE STRATEGY IN MONOCYTIC CELLS
Single Cells
97,1
FSC-H
FSC-A
Live
92,4
FSC-H
Comp-V500-A :: L_D
cd45+
80,0
FSC-H
Comp-V450-A :: CD45
CD11b+F480+
1,20
Comp-PE-A :: F4_80
Comp-APC-Cv7-A :: CD11b
ly6g-
90,3
Comp-FITC-A :: LY6G
Comp-PE-Cv7-A :: CD11c
Control C57BL/6
Infected C57BL/6
Control TNF-R-/-
Infected TNF-R-/-
Ly6c
C
T CD4+
T CD8+
B cell
Monocytic cells
% of Max.
GLUT1
FMO
Isotype control
Uninfected
Infected Pc

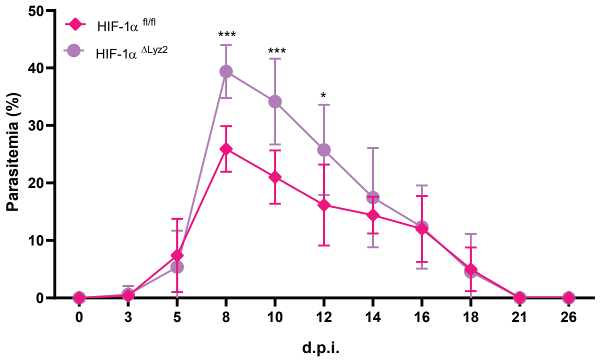

HIF-1α fl/fl
HIF-1α ΔLyz2
***
***
*
Parasitemia (%)
50
40
30
20
10
0
0
3
5
8
10
12
14
16
18
21
26
d.p.i.

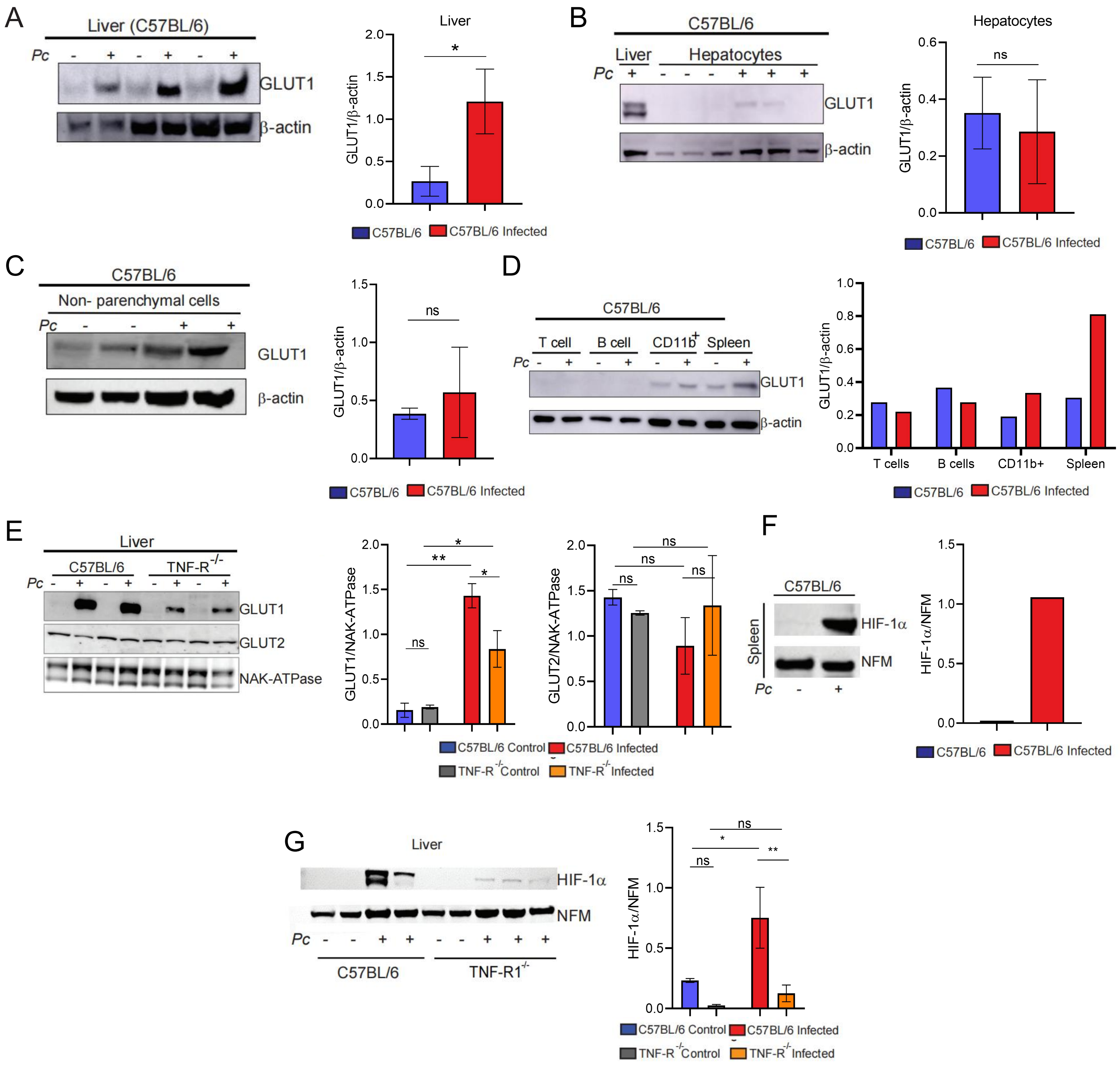
A
Liver (C57BL/6)
Pc
GLUT1
β-actin
Liver
GLUT1/β-actin
C57BL/6
C57BL/6 Infected
B
C57BL/6
Liver
Hepatocytes
GLUT1
β-actin
Hepatocytes
ns
C
C57BL/6
Non- parenchymal cells
D
T cell
B cell
CD11b+
Spleen
T cells
B cells
CD11b+
E
Liver
TNF-R-/-
NAK-ATPase
GLUT2
GLUT1/NAK-ATPase
GLUT2/NAK-ATPase
C57BL/6 Control
TNF-R-/- Control
TNF-R-/- Infected
F
HIF-1α
NFM
HIF-1α/NFM
G
TNF-R1-/-
